# Multiple Pesticides and their Mixtures Tested for Genotoxicity in the Micronucleus Assays on Intestinal Caco-2 Cells

**DOI:** 10.64898/2026.05.16.725095

**Authors:** F. Truzzi, E. Tibaldi, R. Noferini, D. Sgargi, S. Panzacchi, G. Nardali, A. Lorenzini, S. Dilloo, E D’Amen, F Gnudi, G. Dinelli, P.T.J. Scheepers, D Mandrioli

## Abstract

Widespread exposure to multiple pesticides might potentially represent a genotoxic risk to humans. However the effects of these mixtures are largely unknown. Genotoxicity is a key characteristic of carcinogens, and its assessment represents an important component of the overall safety assessment of pesticides. In the present study, *in vitro* micronucleus test on intestinal Caco-2 human cells was performed according to OECD TG 487 in order to ascertain the genotoxicity of ten commonly used pesticides (dose range 0–100 mg L^-1^), tested as individual pesticides or mixtures. Significant dose-related increases in micronuclei were observed for exposures to lambda-cyhalothrin, tebuconazole, glyphosate, deltamethrin, fluopyram and the synergist piperonyl butoxide. Significant increases of micronuclei were also observed at different doses for cypermethrin, acetamiprid and cyprodinil, however these increases were not dose-dependent. Imazalil genotoxicity could not be analyzed due to confounding of high cytotoxicity even at low doses. Results show that the co-formulant piperonyl butoxide was genotoxic to human cell lines at all tested doses. Moreover, glyphosate, acetamiprid and fluopyram showed genotoxic effects at concentrations of 0.01-1.0 mg L^-1^. Although previously reported to be not genotoxic cyprodinil and deltamethrin were observed to be genotoxic to Caco-2 cells. A combination of 3 prioritzided pesticides (acetamiprid, glyphosate, tebuconazole) showed genotoxic effects even at the lowest dose. A combination of 8 prioritized pesticides showed genotoxicity at the highest dose. No synergistic interactions in micronuclei formation were evident in either the mixture of 3 or 8 prioritized pesticides. This study provides important information on the genotoxicity of different widely used pesticides and confirms the validity of a component-based approach in genotoxicity assessment of pesticide mixtures. This study was performed as part of the EU SPRINT (Sustainable Plant Protection Transition: A Global Health Approach) project.

## 1. Introduction

Pesticides are chemical constituents commonly used in farming to prevent or control pests, including insects, rodents, fungi, weeds, and other unwanted organisms [1]. Global pesticide demand has increased substantially over time [2], and the intensive and widespread use of pesticides raises serious environmental and human health concerns [1] [3]. As part of an EU Horizon 2020 funded research project, SPRINT (Sustainable Plant Protection Transition: A Global Health Approach), 209 pesticide residues were examined in study site of 10 separate European countries and 1 in Argentina that included conventional and organic farms. Mixtures of pesticide residues were shown to be omnipresent in agricultural environments (soil, water, sediment, crops, outdoor air) and in farmers’ households (indoor dust samples) [3]. Interestingly, measured concentrations of pesticide residues in soil were also shown to exceed predicted environmental residue concentrations, evidencing that current calculation methods may not reliably estimate the presence of pesticide residues [4]. To date, little is known about health risks posed by these environmentally relevant pesticide mixtures [3].

Acute and chronic health effects of pesticides are serious public health concerns, and within the European context, an urgent need to reduce pesticide use and risk was highlighted by the EU Commission and the European Environment Agency (EEA) due to widespread human exposure [5]. One of the potential chronic side-effects of pesticides is genotoxicity. The strong link between DNA alterations and cancer or chronic degenerative diseases has long been known and widely reported [6], [7] [8]. The general terms ‘genotoxic’ and ‘genotoxicity’ apply to chemical agents which alter the structure, information content, or segregation of DNA, including those which cause DNA damage by interfering with normal replication processes [9]. Hence, genotoxicity assessment represents an important component of the overall safety assessment of pesticides and further investigations are warranting [2]. Genotoxicity is one of the 10 key characteristics of carcinogens considered in IARC hazard assessment. The use of genotoxicity biomarkers, including chromosomal aberrations, micronuclei, sister chromatid exchanges and comets are considered suitable biomarkers for evaluating genotoxicity on cells in *in vitro* testing [10]. Among the methods to test for genotoxicity, the *in vitro* micronucleus assay has gained widespread international regulatory acceptance (OECD [Organization for Economic Co-operation and Development] Test Guideline number 487) and is currently recommended in several authoritative documents [2] [11] [12]. The principle behind the assay is the measurement of micronuclei which are formed during mitosis because of DNA damage from exposure to a test chemical. The cytokinesis-block version (prevention of cytoplasmic division into daughter cells) of the micronucleus assay, is performed by using cytochalasin B (CytoB) which ensures that binucleate cells analyzed have undergone cell division, necessary to ascertain DNA damage and micronucleus formation. Given that the scoring of micronuclei is rendered inaccurate in the presence of cytotoxic doses of pesticides, the measurement of cytotoxicity is a requirement of the OECD Test Guideline Number 487 to ensure that genotoxic assessments are not confounded by cytotoxicity at the applied doses [12].

It is necessary to increase scientific evidence regarding the genotoxicity effects of many pesticides [10], as there is lack of such data and often these are provided by the producers, rather than by independent laboratories [13]. The objective of the present research was to provide genotoxicity assessments of the prioritized (Top 10) pesticides and mixtures of greatest concern based upon SPRINT CSS exposure data. In particular, a quantitative ranking, based on weighted hazard quotient [wHQ] based on SPRINT CSS exposure data and available toxicity data, prioritized the following pesticides: lambda-cyhalothrin (insecticide), cypermethrin (insecticide), deltamethrin (insecticide), tebuconazole (fungicide), glyphosate (herbicide), acetamiprid (insecticide), cyprodinil (fungicide), piperonyl butoxide (synergist and co-formjulant in pyrethroid-based pesticides), fluopyram (fungicide) and imazalil (fungicide). The pesticides were investigated individually as well as in mixtures, namely the Top 3 (acetamiprid + glyphosate + tebuconazole) and the Top 8 (acetamiprid + cypermethrin + cyprodinil + deltamethrin + glyphosate + lambda-cyhalothrin + piperonyl butoxide + tebuconazole), using the international regulatory accepted *in vitro* micronucleus assay on the human intestinal epithelial cell-line, Caco-2. Notably, the genotoxicity of glyphosate has been widely debated [14] [15] [3], while cyprodinil, deltamethrin, fluopyran, and piperonyl butoxide have been generally considered non-genotoxic and scarcely studied in human cells [16] [17] [18] [19] [20] [21, 22]. Furthermore, it was also considered important to extend the dose range to include lower concentrations, even below those expected from exposure to ADI values of the respective pesticides [23] [24], which to date have largely been overlooked in published literature. For this reason, the comparative range selected was from 0.01 mg L^-1^ to 100 mg L^-1^ for each of the pesticides investigated.

## 2. Materials and Methods

Micronucleus assay was performed according to OECD [Organization for Economic Co-operation and Development] Test Guideline number 487 [12].

### 2.1. Materials

The human epithelial colorectal adenocarcinoma Caco-2 cell line (ATCC® HTB-37TM) was purchased distributors in Italy (Sesto San Giovanni, Milan, Italy). Dulbecco’s Modified Eagle Medium (DMEM), fetal bovine serum (FBS), and penicillin-streptomycin (Pen/Strep) were obtained from Gibco (Waltham, MA, USA). Cytochalasin B (CytoB) was from ACROS Organics (Segrate, MI, Italy), Vectashiel Vibrance Antifade Mounting Medium with 4’,6-diamidino-2-phenylindole (DAPI) was purchased from Vector Laboratories (Newark, CA, USA). Imazalil (1-[2-(Allyloxy)-2-(2,4-dichlorophenyl) ethyl] imidazole) (>99% pure), fluopyram (N-[2-[3-Chloro-5-(trifluoromethyl)-2-pyridinyl]ethyl]-2-(trifluoromethyl) benzamide) (>98% pure), piperonyl butoxide (2-(2-Butoxyethoxy) ethyl (6-propylpiperonyl) ether) (>98% pure), cyprodinil (4-Cyclopropyl-6-methyl-N-phenylpyrimidin-2-amine) (> 99% pure), Acetamiprid (N-(6-Chloro-3-pyridylmethyl)-N-cyano-N-methylacetamidine) (>99% pure), glyphosate (N-(Phosphonomethyl) glycine) (>98% pure), tebuconazole (N-(1-(4-Chlorophenyl)-4,4-dimethyl-3-(1H-1,2,4-triazol-1-ylmethyl)-3-pentanol) (>99% pure), deltamethrin (S)-Cyano(3-phenoxyphenyl) methyl (1R,3R)-3-(2,2-dibromoethen-1-yl)-2,2-dimethylcyclopropane-1-carboxylate) (>99% pure), cypermethrin ([Cyano-(3-phenoxyphenyl)methyl]3-(2,2-dichloroethenyl)-2,2-dimethylcyclopropane-1-carboxylate) (>99% pure) and lambda-cyhalothrin (RS)-alpha-cyano-3-phenoxybenzyl 3-(2-chloro-3,3,3-trifluoropropenyl)-2,2,-dimethylcyclopropanecarboxylate) (> 99% pure) were purchased from Merck Life Science S.r.l. (Milan, Italy). All remaining reagents used in the experiments were of analytical grade.

### 2.2. Cell line and culture conditions

The Caco-2 human cell line was used. Cells were cultured in DMEM with 10 % FBS and 1 % Pen/Strep at 37 °C in a humidified incubator with 5 % CO_2_ in tissue culture flasks (75 cm^2^; BD Biosciences, Franklin Lakes, N J, USA). The medium was changed every two days. Prior to experimentation, the cells were trypsinized and cell density evaluated microscopically using a Bürker counting chamber.

### 2.3. Preparation of the individual and combination pesticide treatments

The following ten pesticides were investigated as mono-constituents: acetamiprid, cypermethrin, cyprodinil, deltamethrin, fluopyram, glyphosate, imazalil, lambda-cyhalothrin, piperonyl butoxide and tebuconazole. Two combination pesticide treatments were used, namely the Top 3 (acetamiprid + glyphosate + tebuconazole) and the Top 8 (acetamiprid + cypermethrin + cyprodinil + deltamethrin + glyphosate + lambda-cyhalothrin + piperonyl butoxide + tebuconazole), respectively.

For the individual pesticide treatments, each of the 10 pesticides (5 mg) was-dry mixed with DMEM powder (500 mg) using a ceramic pestle and a mortar. For each pesticide, two separate stock solutions of 100 mg L^-1^ and 10 mg L^-1^ were prepared by diluting the powdered mixtures in distilled water. The 100 mg/L solution was sonicated for 5 min. Prior to use on the cell cultures, all stock solutions were filtered using 0.45 µm diameter polyethersulfone (PES) filters.

For the Top 3, each of acetamiprid, tebuconazole and glyphosate were dry mixed with DMEM powder (1000 mg) using a ceramic pestle and a mortar. A stock solution of 100 mg L^-1^ was prepared following dilution in distilled water. The solution was sonicated for 5 min and filtered with a 0.45 µm diameter PES filter. NaCO_3_ (50 µL 10 mL^-1^) was then added to the solution. Prior to preparing the 10 mg L^-1^ stock solution, three separate 30 mg L^-1^ stock solutions were prepared by dry mixing each pesticide (3 mg) with DMEM powder (1 mg) and adding distilled water. The three 30 mg/L stock solutions were sonicated for 5 min, filtered with 0.45 µm diameter PES filters. NaCO_3_ (50 µL 10 mL^-1^) was added to each solution. Equimolar concentrations of each of the three 30 mg L^-1^ stocks were then combined to produce the 10 mg L^-1^ solution of each pesticide. Similarly, for the Top 8, Acetamiprid, cypermethrin, cyprodinil, deltamethrin, glyphosate, lambda-cyhalothrin, piperonyl butoxide and tebuconazole (10 mg each) were dry mixed with DMEM (1000 mg) and diluted in distilled water to attain a concentration of 100 mg L^-1^. The solution was sonicated for 5 min and then filtered through a 0.45 µm diameter PES filter before the addition of NaCO_3_ (50 µL 10 mL^-1^). As with the Top 3, 80 mg L^-1^ solutions were, similarly, prepared by dry-mixing each pesticide (8 mg) separately in DMEM powder (1 g). The same concentrations of each 80 mg/L pesticide solution were combined to produce the final 10 mg L^-1^ solution.

The 10 mg L^-1^ stock solutions for both the mono-constituent, as well as the Top 3 and Top 8 combination treatments, were subsequently diluted in liquid DMEM to produce concentrations of 0.01, 0.1 and 1 mg L^-1^, respectively.

### 2.3. Culture conditions and administration of pesticide treatments to Caco-2 cells for the measurement of micronuclei

Caco-2 cells were plated in 4-well-chamberslides (3x10^4^ cells in 500ul well^-1^) in Caco-2 medium (DMEM containing with 10 % FBS and 1 % Pen/Strep) and incubated for 24 h. Throughout the procedure, all incubation periods were performed at 37 °C in a humidified incubator with 5 % CO_2._ The Caco-2 medium was removed and replaced with pesticide treatments (10 mono-constituent treatments and two combinations) at concentrations of 0.01, 0.1, 1.0, 10 and 100 mg L^-1^, respectively. The negative untreated control (CTRL) contained only DMEM. The experimental exposure period was 4 h. Thereafter, the medium containing the treatments was removed and replaced with fresh Caco-2 medium. The plates were incubated for a further 48 h, after which 4 μg mL^-1^ DMEM CytoB was added to each treatment well. The CTRL contained only DMEM. The plates were incubated for 24 h. Thereafter, the DMEM was aspirated, and the cells washed with 1 x PBS (500 µL well^-1^), followed by 75 mM KCL (500 µL well^-1^) for 2 min at room temperature. The KCL was then removed and plates washed again with 1 x PBS (500 µL well^-1^). Cells were subsequently fixed in a solution of methanol: glacial acetic acid (3:1) and incubated for 10 min at room temperature. The solution was then removed. After removing the chamber side walls, the cell monolayers were either stained immediately for use or maintained at -20 °C until a suitable time.

### 2.5. Staining and microscopic viewing of micronuclei in Caco-2 cells

Unstained slides maintained at -20 °C were removed and allowed to thaw at room temperature for 10 min. Thereafter, slides were incubated in PBS in a vertical position for 10 min, then placed in a horizontal position and stained. Staining was performed by adding Vectashield Vibrance Antifade Mounting Medium with DAPI. Since each slide contained 4 treatment wells, one drop of DAPI was added to each treatment. A cover slip was placed over the slide and sealed with transparent nail polish. Slides were viewed using a fluorescent NIKON Ni-E microscope (NIKON, Japan). To take photos, the image software used was NIS ELEMENTS AR Version 5.30.06 64 bit (NIKON, Japan). For imaging, the monochromatic camera (Qi2) was first chosen to activate the program. Thereafter, the blue filter for DAPI staining was selected with the selected Eyepiece-EPI to view the cells at a magnification of x 40. After determining the location of the cells, the magnification objective was changed to x 60 magnification and immersion oil added. The cells were viewed from the computer by replacing the Eyepiece-EPI selection to Qi2-EPI. A total of 40 photos were taken for each slide (10 photos well^-1^) at random positions, and each photo saved in jpeg format for analysis and elaboration of digital images.

### 2.6. Photographic scoring criteria of Caco-2 cells for the CBPI index and the presence of micronuclei

Since CytoB was used, estimates of cytotoxicity were based on the Cytokinesis-Block Proliferation Index (CBPI). The CBPI scoring procedure includes the number of mononucleated, binucleated and polynucleated cells, respectively, as well as the total number of cells. Additionally, at least 1000 binucleated cells/culture were examined for the presence of micronuclei. The number of binucleated cells in the positive CTRLs and treated groups subject to high pesticide dose (high toxicity index) may be < 1000 cells. Binucleated cells were scored based on the following criteria: the presence of nuclei that were separate and of equal size, the presence of nuclei that may be in contact but with distinguishable nuclear boundaries, and nuclei that were linked by nucleoplasmic bridges. Exclusion criteria included cells in which the main nuclei were undergoing apoptosis (micronuclei may have already disappeared due to apoptosis). The inclusion of mononucleates in the calculation accounts for cytostasis and as such CBPI provides an estimate of cell proliferation. CBPI for the pesticide-treated cells and CTRLs were calculated separately.

The following formula was used:

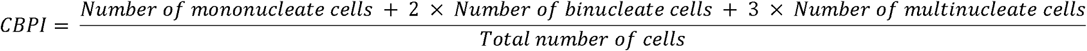

The cytotoxicity estimate was calculated as follows:

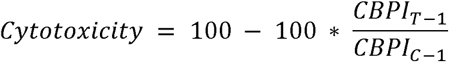

Where T = test pesticide treated cultures, C = CTRL cultures

The criteria for scoring micronuclei were as follows: diameters not exceeding one third of the main the main nucleus, the presence of micronuclei separate from the main nucleus, the presence of micronuclei that marginally overlap with the main nucleus but with distinguishable nuclear boundaries, a similar staining pattern between micronuclei and the main nucleus, and finally that micronuclei be located near the edge of the main nucleus of the binucleated cell.

### 2.7. Analysis of images

Digital images were analyzed using FIJI Image J-win 64 which was downloaded free of charge. The jpeg images were opened and the following procedure sequence followed: Plugins-Analysed-Cell Counter-Initialized. The types of cells that were counted included mononucleated, binucleated and polynucleated cells, respectively, as well as those containing micronuclei. The software recorded the total number of each cell type counted. Numbers were assigned to distinguish the cell types as follows: binucleated cells (1), polynucleated cells (2), mononucleated cells (3) and micronuclei (4). The data obtained from the counting with the Image J software was transferred to an Excel file.

### 2.8. Statistical analysis

Mononucleated, binucleated, multinucleated cells, and the number of micronuclei was assessed for samples from all groups. The number of micronuclei on the total of binucleated cells was compared between the controls (treated with CytoB) and all doses of each treatment using a 1-sided Fisher Exact test. Furthermore, the presence of a linear trend across doses was verified using a Cochran-Armitage trend test. Results were considered statistically significant when the p-values were < 0.05. Statistical analyses were performed using the German Cancer Research Center software (https://biostatistics.dkfz.de/) and Stata 18 software (StataCorp 2023. Stata Statistical Software: Release 18. College Station, TX: StataCorp LLC).

### 2.9. Validation of the efficacy of the negative and positive controls for the in vitro micronucleus test

In the present study, the negative CTRL, treated with CytoB only, was first tested to confirm the formation of binucleate cells and to provide the basal estimate of micronuclei in a minimum number of 1000 binucleate cells. A total of 34 micronuclei were counted for a total of 1037 Caco-2 binucleate cells (Table 1). Then, Mito C, known to induce chromosomal breakage, was the selected clastogenic positive CTRL. Efficacy of the positive CTRL was verified from the formation of 217 micronuclei in 1009 Caco-2 binucleate cells (Table 1). Neither CytoB nor Mito C were shown to be cytotoxic to the Caco-2 cells at that concentration. Cytotoxicity ranged from 0.96 to 1.90 % (Table 1). These values were significantly below the 55.0 ± 5.0 % set target, indicating that the administration of the positive and negative CTRLs did not interfere with the scoring of micronuclei due to cytotoxic effects on the cells.

**Table 1.** The effects of the non-pesticide containing negative control (CTRL) with Cytochalasin B (CytoB) and positive CTRL with clastogenic mitomycin C (Mito C) on the number of binucleated cells and genotoxic micronuclei as well as the cytotoxicity percentages in Caco-2 cells.

|  | Pesticide<br>Dose | Binucleated<br>cells (number) | Micronuclei<br>(number) | Significant<br>increase | Cytotoxicity<br>(%) |
| --- | --- | --- | --- | --- | --- |
| CTRL (only CytoB) | 0 | 1027 | 34 |  | 0.96 |
| Mito C | 0 | 1009 | 217 | * | 1.90 |

\*1-sided Fisher Exact test (P<0.01)

## 3. Results

The results of the micronuclei genotoxicity assays of each of the 10 single pesticides and the 2 mixtures tested at 5 dose levels are provided here. Pairwise comparisons of the incidence of micronuclei over binucleated cells were performed within the dose range of 0 – 100 mg L^-1^ for the each of the pesticides as mono-constituents (Figs. 1-9, Table 2 and Table 3) and in the combination treatments (Figs. 10-11). Significantly increasing concentrations of pesticides with the highest incidence of micronuclei present at the final dose of 100 mg L^-1^ were evident from exposure to lambda-cyhalothrin (Fig. 1), deltamethrin (Fig. 3), tebuconazole (Fig. 4), glyphosate (Fig. 5), piperonyl butoxide (Fig. 8), fluopyram (Fig. 9) and the Top 8 (Fig.11), respectively. The Cochran-Armitage trend test showed a significant dose-related increase for lambda-cyhalothrin deltamethrin, tebuconazole, glyphosate, piperonyl butoxide, fluopyram and the Top 8. A total of 273 micronuclei were counted at 100 mg L^-1^ glyphosate, which exceeded the number induced by the known clastogenic agent, mitomycin C (Fig. 5, Table 1). At 100 mg L^-1^, the numbers of micronuclei for fluopyram were 150, with numbers declining close to 100 for tebuconazole and piperonyl butoxide before decreasing to approximately 50 micronuclei for deltamethrin, lambda-cyhalothrin and Top 8, respectively.

**Fig. 1.**
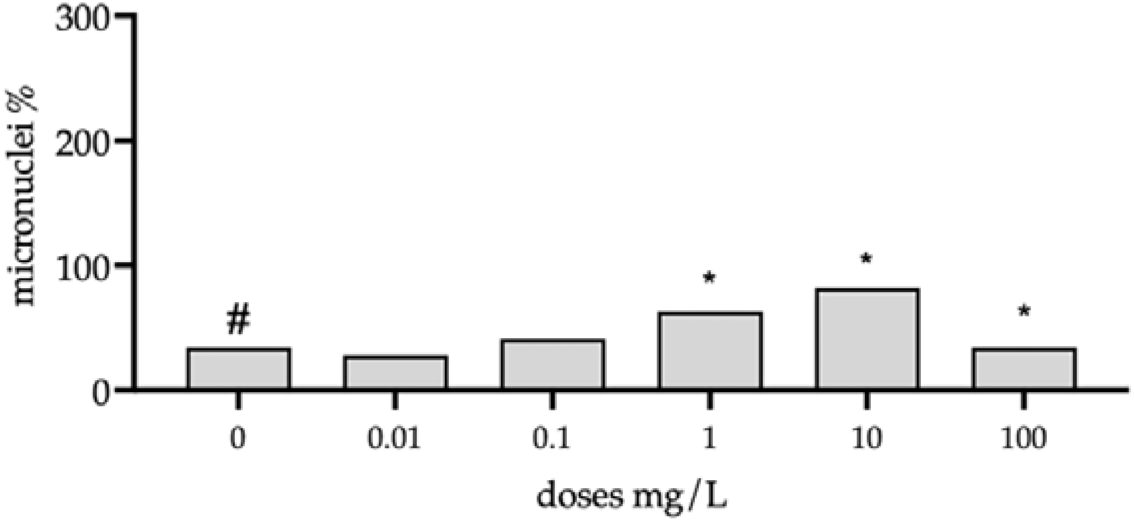
Effects of pure lambda-cyhalothrin, within the dose range 0 – 100 mg L^-1^, on the number of micronuclei per 1000 binucleate Caco-2 cells. Data for each dose is expressed as a single value. Pairwise comparisons of the incidence of micronuclei over binucleated cells were performed using the 1-sided Fisher’s exact test (*P < 0.05). The presence of increased linear trend was evaluated using one sided Cochran-Armitage trend test (P < 0.05).

**Fig. 2.**
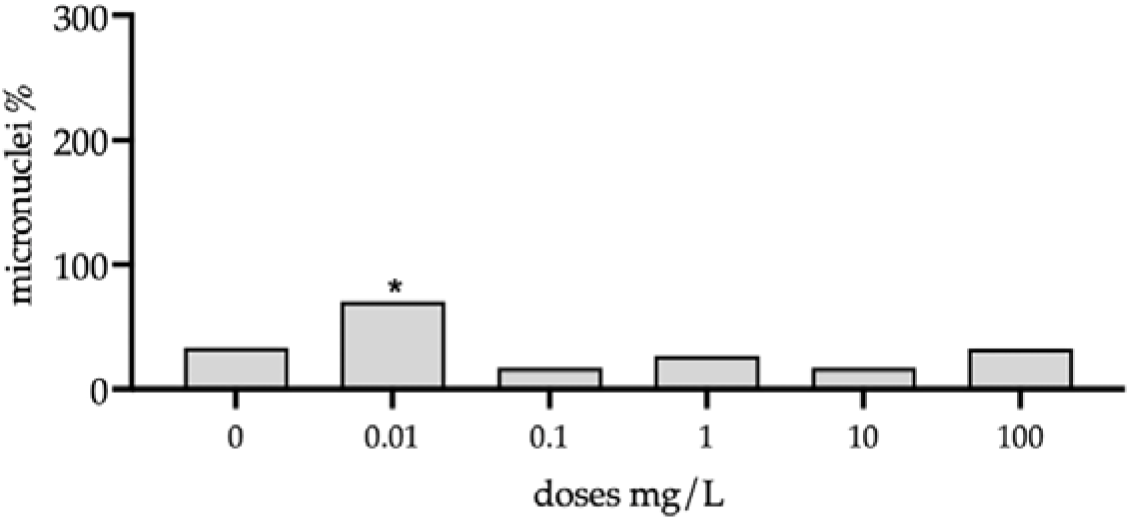
Effects of pure cypermethrin, within the dose range 0 – 100 mg L^-1^, on the number of micronuclei per 1000 binucleate Caco-2 cells. Data for each dose is expressed as a single value. Pairwise comparisons of the incidence of micronuclei over binucleated cells were performed using the 1-sided Fisher’s exact test (*P < 0.05). The presence of increased linear trend was evaluated using one sided Cochran-Armitage trend test (P < 0.05).

**Fig. 3.**
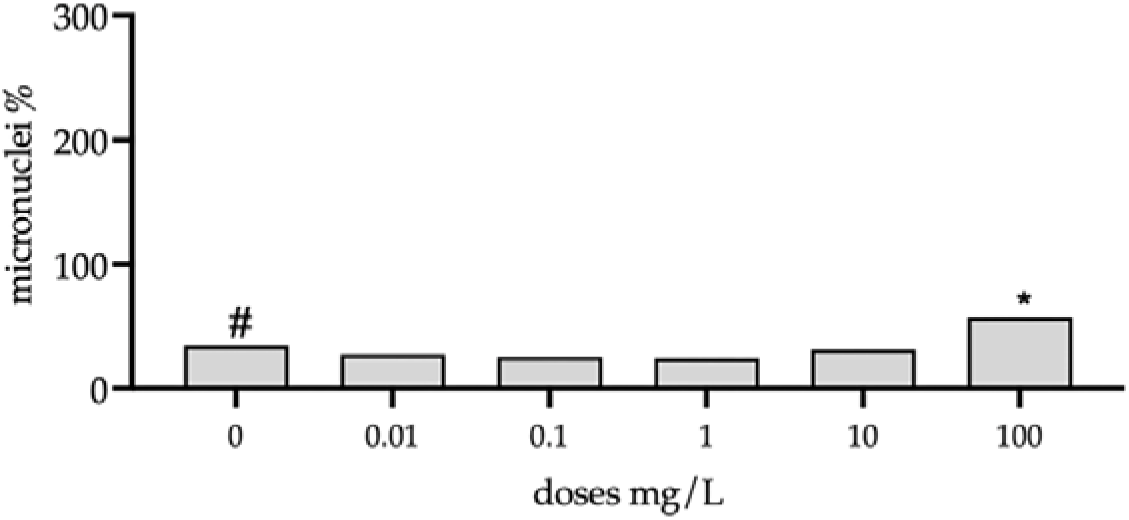
Effects of pure deltamethrin, within the dose range 0 – 100 mg L^-1^, on the number of micronuclei per 1000 binucleate Caco-2 cells. Data for each dose is expressed as a single value. Pairwise comparisons of the incidence of micronuclei over binucleated cells were performed using the 1-sided Fisher’s exact test test (*P < 0.05). The presence of increased linear trend was evaluated using one sided Cochran-Armitage trend test (P < 0.05).

**Fig. 4.**
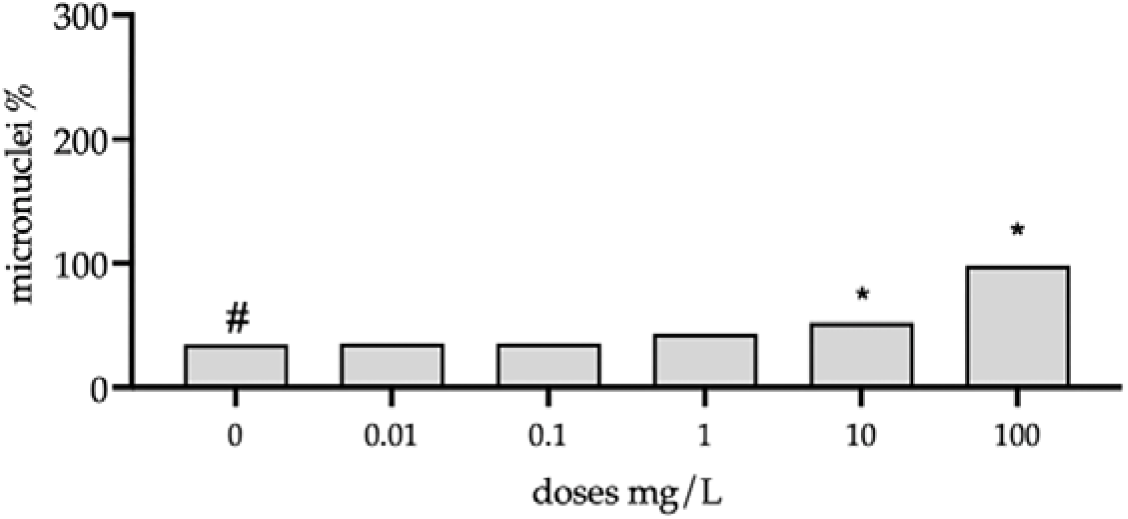
Effects of pure tebuconazole, within the dose range 0 – 100 mg L^-1^, on the number of micronuclei per 1000 binucleate Caco-2 cells. Data for each dose is expressed as a single value. Pairwise comparisons of the incidence of micronuclei over binucleated cells were performed using the 1-sided Fisher’s exact test (*P < 0.05). The presence of increased linear trend was evaluated using one sided Cochran-Armitage trend test (P < 0.05).

**Fig. 5.**
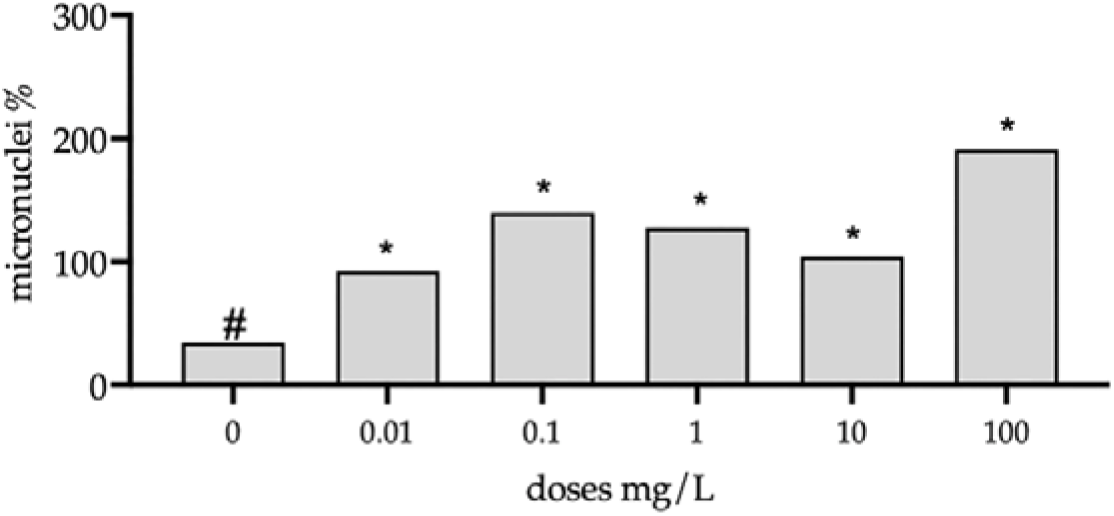
Effects of pure glyphosate, within the dose range 0 – 100 mg L^-1^, on the number of micronuclei per 1000 binucleate Caco-2 cells. Data for each dose is expressed as a single value. Pairwise comparisons of the incidence of micronuclei over binucleated cells were performed using the 1-sided Fisher’s exact test (*P < 0.05). The presence of increased linear trend was evaluated using one sided Cochran-Armitage trend test (P < 0.05).

**Fig. 6.**
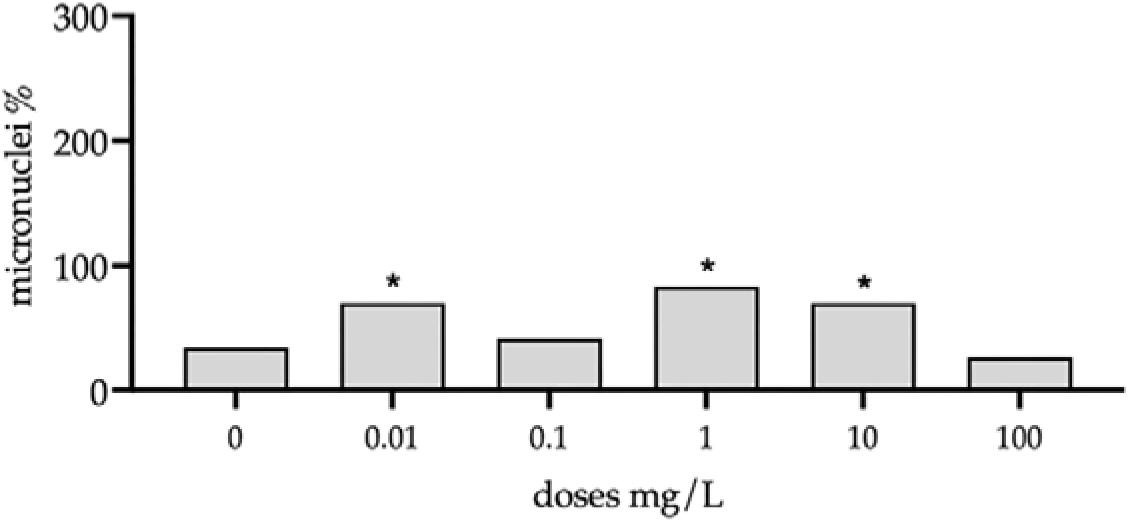
Effects of pure acetamiprid, within the dose range 0 – 100 mg L^-1^, on the number of micronuclei per 1000 binucleate Caco-2 cells. Data for each dose is expressed as a single value. Pairwise comparisons of the incidence of micronuclei over binucleated cells were performed using the 1-sided Fisher’s exact test (*P < 0.05). The presence of increased linear trend was evaluated using one sided Cochran-Armitage trend test (P < 0.05).

**Fig. 7.**
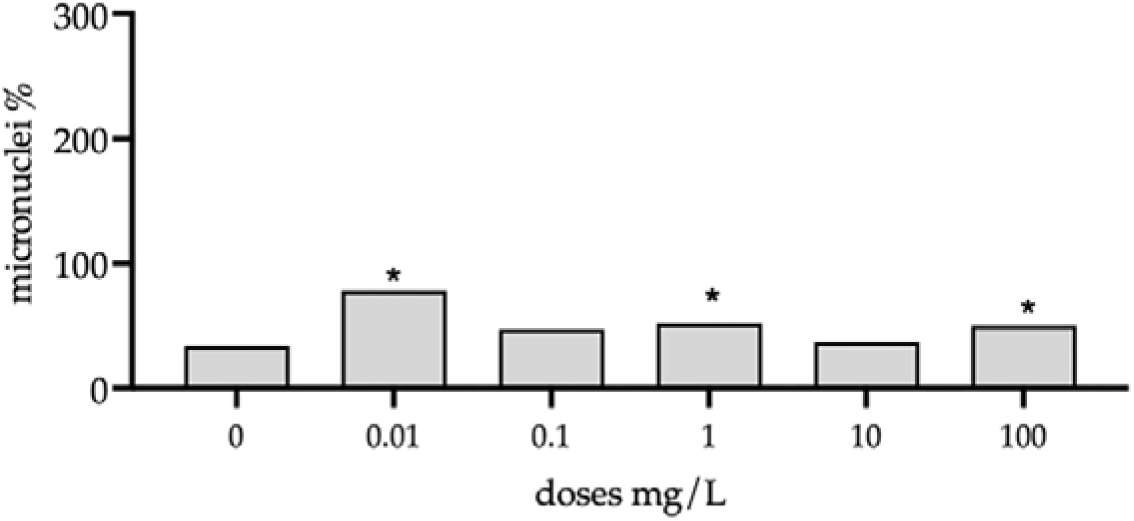
Effects of pure cyprodinil, within the dose range 0 – 100 mg L^-1^, on the number of micronuclei per 1000 binucleate Caco-2 cells. Data for each dose is expressed as a single value. Pairwise comparisons of the incidence of micronuclei over binucleated cells were performed using the 1-sided Fisher’s exact test (*P < 0.05). The presence of increased linear trend was evaluated using one sided Cochran-Armitage trend test (P < 0.05).

**Fig. 8.**
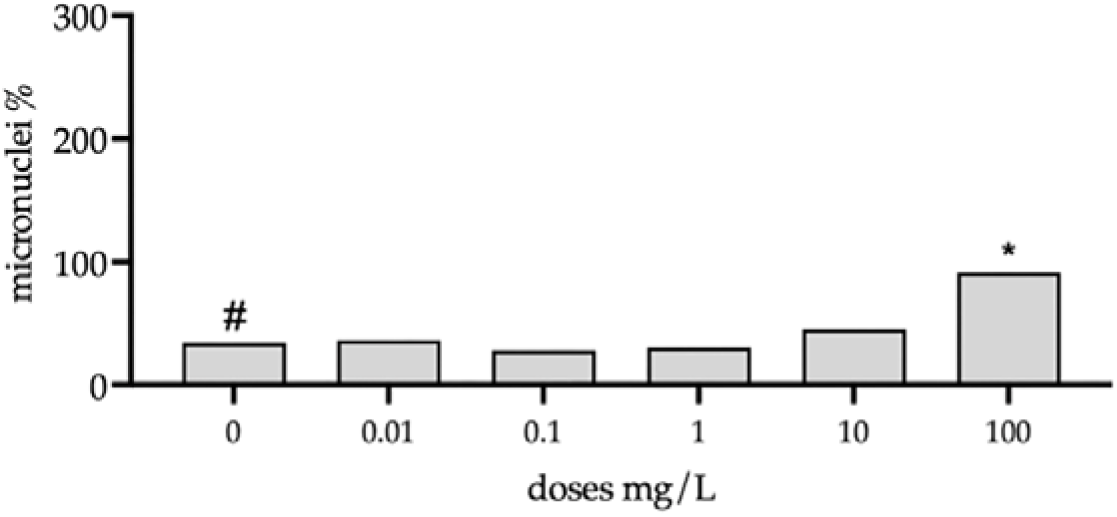
Effects of pure piperonyl butoxide, within the dose range 0 – 100 mg L^-1^, on the number of micronuclei per 1000 binucleate Caco-2 cells. Data for each dose is expressed as a single value. Pairwise comparisons of the incidence of micronuclei over binucleated cells were performed using the 1-sided Fisher’s exact test (*P < 0.05). The presence of increased linear trend was evaluated using one sided Cochran-Armitage trend test (P < 0.05).

**Fig. 9.**
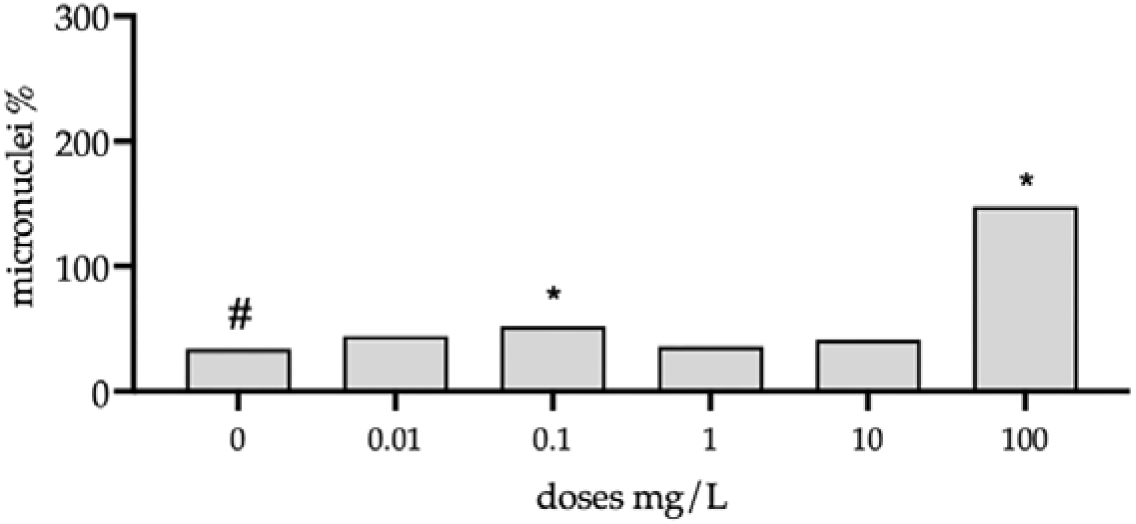
Effects of pure fluopyram, within the dose range 0 – 100 mg L^-1^, on the number of micronuclei per 1000 binucleate Caco-2 cells. Data for each dose is expressed as a single value. Pairwise comparisons of the incidence of micronuclei over binucleated cells were performed using the 1-sided Fisher’s exact test (*P < 0.05). The presence of increased linear trend was evaluated using one sided Cochran-Armitage trend test (P < 0.05).

**Fig. 10.**
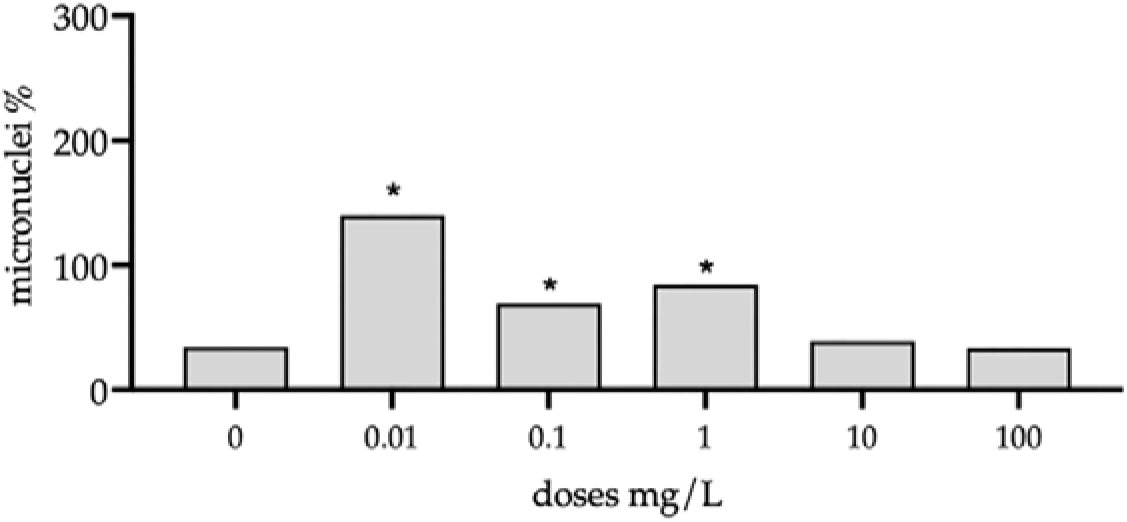
Effects of a combination mixture of pesticides called Top 3 (pure acetamiprid, tebuconazole, glyphosate), within the dose range 0 – 100 mg L^-1^, on the number of micronuclei per 1000 binucleate Caco-2 cells. Data for each dose is expressed as a single value. Pairwise comparisons of the incidence of micronuclei over binucleated cells were performed using the 1-sided Fisher’s exact test (*P < 0.05). The presence of increased linear trend was evaluated using one sided Cochran-Armitage trend test (P < 0.05).

**Fig. 11.**
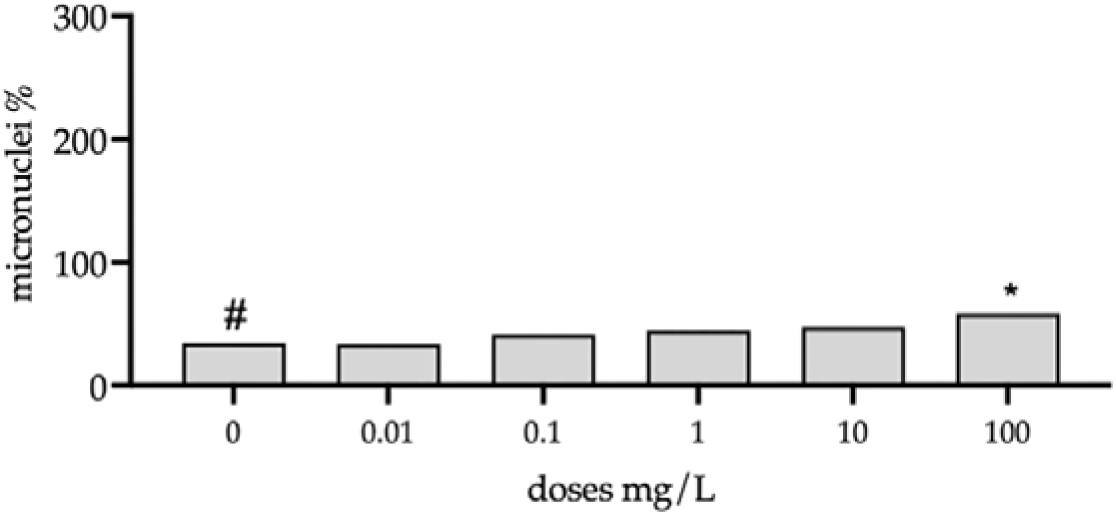
Effects of a combination mixture of pesticides called Top 8 (pure lambda-cyhalothrin, cypermethrin, deltamethrin, tebuconazole, glyphosate, acetamiprid, cyprodinil, piperonyl butoxide), within the dose range 0 – 100 mg L^-1^, on the number of micronuclei per 1000 binucleate Caco-2 cells. Data for each dose is expressed as a single value. Pairwise comparisons of the incidence of micronuclei over binucleated cells were performed using the 1-sided Fisher’s exact test (*P < 0.05). The presence of increased linear trend was evaluated using one sided Cochran-Armitage trend test (P < 0.05).

**Table 2.** The effect of imazalil doses on binucleate cell number, micronuclei number and cytotoxicity in Caco-2 cells.

| Dose (mg/L) | Binucleated cells (number) | Micronuclei (number) | Cytotoxicity (%) |
| --- | --- | --- | --- |
| 0 | 1027 | 34 |  |
| 0.01 | 1008 | 14 | 38.82 |
| 0.1 | 1024 | 12 | 39.34 |
| 1 | 1010 | 13 | 38.66 |
| 10 | 1011 | 12 | 42.00 |
| 100 | 128 | 13 | 82.61 |

**Table 3.** Cytotoxicity assessments on Caco-2 cells for the pesticides, as mono-constituents, for all doses (0 – 100 mg L^-1^).

| **Substance** | **Dose** | **% Cytotoxicity** |
| --- | --- | --- |
| Lambda-cyhalothrin | 0.01 | 42.08 |
| Lambda-cyhalothrin | 0.1 | 39.43 |
| Lambda-cyhalothrin | 1 | 37.44 |
| Lambda-cyhalothrin | 10 | 39.91 |
| Lambda-cyhalothrin | 100 | 57.54 |
| Cypermethrin | 0.01 | 26.56 |
| Cypermethrin | 0.1 | 26.52 |
| Cypermethrin | 1 | 32.02 |
| Cypermethrin | 10 | 28.72 |
| Cypermethrin | 100 | 12.67 |
| Deltametrin | 0.01 | 36.63 |
| Deltametrin | 0.1 | 34.75 |
| Deltametrin | 1 | 40.05 |
| Deltametrin | 10 | 38.85 |
| Deltametrin | 100 | 38.22 |
| Tebuconazole | 0.01 | 36.88 |
| Tebuconazole | 0.1 | 37.55 |
| Tebuconazole | 1 | 36.11 |
| Tebuconazole | 10 | 38.00 |
| Tebuconazole | 100 | 34.92 |
| Glyphosate | 0.01 | 24.17 |
| Glyphosate | 0.1 | 10.40 |
| Glyphosate | 1 | 11.71 |
| Glyphosate | 10 | 10.78 |
| Glyphosate | 100 | 22.98 |
| Acetamiprid | 0.01 | 19.78 |
| Acetamiprid | 0.1 | 27.47 |
| Acetamiprid | 1 | 18.92 |
| Acetamiprid | 10 | 20.00 |
| Acetamiprid | 100 | 31.72 |
| Cypronidil | 0.01 | 35.26 |
| Cypronidil | 0.1 | 38.62 |
| Cypronidil | 1 | 40.43 |
| Cypronidil | 10 | 42.88 |
| Cypronidil | 100 | 42.96 |
| Piperonyl butoxide | 0.01 | 36.78 |
| Piperonyl butoxide | 0.1 | 36.22 |
| Piperonyl butoxide | 1 | 34.54 |
| Piperonyl butoxide | 10 | 34.87 |
| Piperonyl butoxide | 100 | 36.25 |
| Fluopyran | 0.01 | 35.25 |
| Fluopyran | 0.1 | 34.48 |
| Fluopyran | 1 | 36.34 |
| Fluopyran | 10 | 38.99 |
| Fluopyran | 100 | 13.55 |

**Table 4.** Cytotoxicity assessments on Caco-2 cells for the pesticides in combination for all doses (0 – 100 mg L^-1^).

| Substance | Dose | % Cytotoxicity |
| --- | --- | --- |
| Top3 | 0.01 | 36.63 |
| Top3 | 0.1 | 39.72 |
| Top3 | 1 | 39.57 |
| Top3 | 10 | 37.41 |
| Top3 | 100 | 38.24 |
| Top8 | 0.01 | 40.54 |
| Top8 | 0.1 | 43.97 |
| Top8 | 1 | 44.53 |
| Top8 | 10 | 44.80 |
| Top8 | 100 | 42.10 |

A significantly higher number of micronuclei was evident over all doses of glyphosate (Fig. 5), showing that this pesticide was the most genotoxic to Caco-2 cells in comparison the remaining mono-constituent and combination pesticides. Of interest, the presence of micronuclear doses in and around the ADI values (0. 5 mg kg^-1^ body weight) was also evident for glyphosate (Fig. 5).

The non-dose-related increases in micronuclei formation were characteristic for cypermethrin (Fig. 2), acetamiprid (Fig. 6), cyprodinil (Fig. 7) and the Top 3 (Fig. 10). For cypermethrin, cyprodinil and the Top 3, respectively, the highest micronuclei formation occurred at the lowest concentration of 0.01 mg L^-1^ compared to all other doses. Micronuclei formation was only evident at 0.01, mg L^-1^ for cypermethrin (Fig. 2), between 0.01-1 mg L^-1^ for the Top 3 (Fig. 11), between 0.01-10 mg L^-1^ for acetamiprid (Fig. 6), and between 0.01-100 mg L^-1^ for cyprodinil (Fig. 7), respectively. These values coincide with those in the range around the respective ADI values.

Imazalil could not be effectively assessed for genotoxic potential based on cytotoxicity effects observed at the 100 mg L^-1^ dose, coinciding with a significant decrease in the number of binucleate cells (Table 2). Moreover, the number of micronuclei at all doses were shown to be inferior to the baseline number of micronuclei in the CytoB CTRL (Table 1, Table 2).

### 3.2.1 Comparison of the combination pesticides in relation to the respective individual mono-constituents on micronuclei formation in binucleate Caco-2 cells

For the Top3 (acetamiprid + glyphosate + tebuconazole), both glyphosate and tebuconazole demonstrated dose-related increases in micronuclei formation as mono-constituents whilst acetamiprid was non-dose-related (Figs. 4-6). In combination, the significantly highest genotoxic effect at the lowest concentration at 0.01 mg L^-1^ appeared additive from the singular contributions of acetamiprid and glyphosate (Fig. 10). No effect was evident at the 10-100 mg L^-1^ concentration range (Fig. 10), even though the highest micronuclei formation was recorded at this range for both glyphosate and tebuconazole individually (Figs. 4-5).

For the Top 8 (acetamiprid + cypermethrin + cyprodinil + deltamethrin + glyphosate + lambda-cyhalothrin + piperonyl butoxide + tebuconazole), individually, five pesticides displayed dose-related genotoxic increases (Figs. 1, 3-5, 8), whereas three displayed non-dose-related increases (Figs. 2, 6-7). In combination, only the 100 mg L^-1^ dose produced significantly higher numbers of micronuclei (Fig. 11), which were equivalent in number to that shown individually for lambda-cyhalothrin (Fig. 1), deltamethrin (Fig. 3) and cyprodinil (Fig. 7), respectively. No synergistic interactions were reported for either the Top 3 or the Top 8.

### 3.3. Effects of the doses of the ten pesticides and two combination treatments on the cytotoxicity in Caco-2 cells

For the cytotoxicity assessment, the CBPI was calculated for each pesticide dose, a requirement when evaluating genotoxic effects as outlined in the OECD Test Guideline No. 487 (OECD, 2023). A 55 ± 0.5 % cytotoxicity target was provided (OECD, 2023). Micronuclei frequencies can be significantly over or alternatively underestimated at doses that induce toxicity more than 55 %, whereas all values below 55 % are not considered cytotoxic. From the present results, the range of doses (0.01-100 mg L^-1^) used for all the pesticides analyzed were not cytotoxic to the accurate estimation of micronuclei (Table 3 and 4). The only exception was for Imazalil at 100 mg L^-1^ (Table 2) which was then not considered in the analysis.

## 4. Discussion

Given the widespread public health concerns from the use of potentially genotoxic pesticides in the agricultural sector, the present research addressed the following critical points : (1) reducing the use of animals in risk assessments; (2) use of new *in vitro* approaches that can more efficiently test, classify, and prioritize chemicals for evaluation and regulation; (3) the use of human cells in *in vitro* approaches, more specifically the use of epithelial cells as a “site of first contact” for which is there is demand; (4) strict adherence to internationally recognized protocols for comparative purposes when comparing research; (5) selection of suitable pesticide doses in which genotoxic assessments were not confounded by cytotoxicity of the applied doses, and (6) re-evaluation of known pesticides for which research on human cells was either lacking or produced inconsistent or confounding results.

In response to points 1-3 above, the present study employed an *in vitro* approach, specifically using human intestinal epithelial Caco-2 cells as the “site of first contact via oral ingestion” model system. To the best of our knowledge, only one study using Caco-2 cells was reported the genotoxicity of acetamiprid, but cytotoxicity was also a confounding factor at certain doses in the assay system [25].

In line with points (4) and (5), the present study strictly adhered to the recommendations of the *in vitro* micronucleus test [12], which has gained increasingly widespread international regulatory acceptance for *in vitro* genotoxic testing. From the controls implemented (presence of the required number of binuclear cells at all doses, absence of interfering cytotoxicity at the selected pesticide doses, the presence of numerous micronuclei when using a known clastogenic agent), the efficacy of the *in vitro* micronucleus test procedure for Caco-2 cells was verified. This study addressed point (6) by reassessing the genotoxic effects in human cell lines of the ten pesticides prioritized by the SPRINT project as being of concern for EU farmers and consumers.

Overall, research on genotoxic effects of known pesticides on human cells is scarce [11]. There are either no or insufficient studies using human cells for piperonyl butoxide, cyprodinil, deltamethrin and fluopyran, which are largely considered non-genotoxic. Nonetheless, of particular interest is the considerable controversy regarding the genotoxicity of glyphosate [14] [15] [3], despite IARC in 2015 state that “there is strong evidence that glyphosate causes genotoxicity”. In the present study, glyphosate exposure was shown to induce the highest percentage formation of micronuclei compared to the remaining pesticides in mono-constituent form. Glyphosate was shown to be genotoxic to epithelial Caco-2 cells from 0.01 mg L^-1^ to 100 mg L^-1^ (ca 0.06-600 µM), contrasting with previous reports using human lymphocyte cells. In white blood cells, it was shown that glyphosate alone was only genotoxic at concentrations exceeding either 100 µM [26] [15] [3] or 200 µM [14] and 300 µM [27]. In TK6 cells, genotoxicity was only shown above 10 mM glyphosate [28]. To the best of our knowledge, this is the first report using human epithelial cells showing genotoxicity by glyphosate from a single exposure alone at low concentration ranges of 0.01-1.0 mg L^-1^. Given that life-time daily tolerance to glyphosate without appreciable health concerns is set at 0.1 and 0.5 mg/kg body weight for accepted operator exposure levels (AOEL) and consumers (ADI), respectively [23] [24], the study highlights potential genotoxic effects to human epithelial cells from a single exposure at levels that are inferior and equal to current AOEL and ADI values.

The present study showed for the first time that the synergist and co-formulant piperonyl butoxide was genotoxic to human cells. Piperonyl butoxide was genotoxic to Caco-2 cells at 100 mg L^-1^, despite being previously considered a non-genotoxic compound [18]. More recently, the need to reassess the hazards of piperonyl butoxide was highlighted [20]. In the latter editorial, genotoxic effects on animals were mentioned with no studies on human cells being reported. Moreover, although well-recognized to be highly cytotoxic compound, the administration of cyprodinil to Caco-2 cells, at a concentration as low as 0.01 mg L^-1^, resulted in micronuclei formation from a single exposure, despite being considered non-genotoxic from previous studies conducted on white blood cells [17] [21]. As with glyphosate, the 0.03 mg kg^-1^ ADI of cyprodinil [23] [24] is equivalent to the value inducing genotoxic effects in Caco-2 from a single dose.

Furthermore, micronuclei formation was shown for the first time in human cells following exposure to deltamethrin alone, despite being considered non-genotoxic in previous reports [29] [16] [19]. Present results corroborated the genotoxicity potential of fluopyran (0.1, 100 mg L^-1^) which was only recently shown to be genotoxic for the first time in blood lymphocytes, measured at concentrations exceeding 50 mg L^-1^ [30]. Lambda-cyhalothrin, cypermethrin, tebuconazole, acetamiprid were, similarly, all genotoxic to human intestinal Caco-2 cells. Due to cytotoxicity induced by imizalil, this was the only pesticide that could not be effectively evaluated for genotoxicity in the present study.

Present results corroborated the presence of genotoxicity at doses within the range used in the present study on cell lines for lambda-cyhalothrin [31] [32], cypermethrin [33] [34] [35], tebuconazole [36] and acetamiprid [37] [25], respectively. However, for acetamiprid, micronuclei formation in Caco-2 cells was also observed between 0.01-10 mg L^-1^, a range either not investigated in previous studies [37] [25] or investigated but found to be negative for increase of micronuclei in white blood cells [38]. Similarly, micronuclei formation at 0.01 mg L^-1^ cypermethrin in the present study corroborated previous results for this compound at 0.025 mg L^-1^ [35], highlighting the importance of administering pesticides at ADI levels. Previously, cypermethrin was only shown to be genotoxic at higher concentrations, although the 0.01 – 5 mg L^-1^ range was not investigated [33] [34].

For the all the pesticides analyzed in mono-constituent form, two response mechanisms were evident, namely a dose-dependent increase in micronuclei formation and a non-dose-dependent incidence of micronuclei. Regarding the dose-dependent increases between 0 – 100 mg L^-1^, lambda-cyhalothrin, deltamethrin, tebuconazole, glyphosate, piperonyl butoxide and fluopyram displayed a dose-dependent increase in micronuclei formation in Caco-2 cells. Previous work demonstrating dose-dependent increases within regions of the concentration ranges examined were evident for lambda-cyhalothrin [31], tebuconazole [36], glyphosate [26] [27], and fluopyram [30]. Regarding non-dose dependent responses, cypermethrin, acetamiprid and cyprodinil displayed non-dose-dependent incidences in micronuclei.

For the first time, the present study also addressed pesticides that have either not yet been tested (Top 3 and Top 8), or for which there is no previously reported evidence of adequate testing on human cell lines. Synergistic effects of pesticides in combination have been widely reported in the literature, and it was considered important to investigate this possibility. However, no synergistic interactions in micronuclei formation were evident in either the Top 3 or Top 8. Of relevance is that in the combination of the Top 3 (acetamiprid + glyphosate + tebuconazole), interactions between the respective constituents resulted in significant micronuclei formation at the lowest doses, not evident for either glyphosate or tebuconazole singularly, which were dose dependent. To the best of our knowledge there are no reports of acetamiprid + glyphosate + tebuconazole combinations in the available literature presenting results that can be compared to the present results. For the Top 8, genotoxicity was evident at the highest dose, presumably because five of the eight pesticide constituents demonstrated dose-dependent increases in micronuclei.

## 5. Conclusions

In conclusion, the present study showed that lambda-cyhalothrin, cypermethrin, deltamethrin, tebuconazole, glyphosate, cyprodinil, piperonyl butoxide, fluopyram, and two mixtures of 3 and of 8 aforementioned compounds were genotoxic to intestinal epithelial Caco-2 cells. Dose-dependent responses in micronuclei formation were evident for lambda-cyhalothrin, tebuconazole, glyphosate, deltamethrin, fluopyram and piperonyl butoxide. Instead, non-dose-dependent increases in micronuclei were evident for cypermethrin, acetamiprid and cyprodinil. Glyphosate by single testing andcombinedwith acetamiprid and fluopyram showed significant micronuclei increases in intestinal cells at lower doses (0.01-0.1 mg L^-1^) than those reported previously for human white blood cells. These results highlighted importance of extending the tested dose-ranges to include ADI values. Like the results shown previously for cypermethrin on blood lymphocytes and a HepG2 cell line [35], present results also indicated genotoxicity in the same range as the ADI values for this compound. Whether intestinal epithelial cells are more sensitive to pesticides compared to white blood cells cannot be ascertained based on the overall scarcity of comparagel studiers in human cell lines. Noteworthy, the present study showed for the first time that the synergist and co-formulant piperonyl butoxide in pyrethroid-based products was observed to be genotoxic to human cell lines. Moreover, despite being considered non-genotoxic, both cyprodinil and deltamethrin were shown for the first time to be genotoxic to Caco-2 cells. This preliminary study was successful in increasing the scientific evidence regarding the genotoxicity effects of commonly used pesticides, highlighting the need for further research on human cells and incorporating lower-dose concentration ranges in risk assessment studies. This study also confirms the validity of a component-based approach in genotoxicity assessment of pesticide mixtures.

## CRediT authorship contribution statement

**FS:** conceptualization, data curation and formal analysis, investigation, visualization, writing the original work, reviewing and editing

**ET:** conceptualization, data curation and formal analysis, investigation, visualization, reviewing and editing

**RN**: investigation

**DS:** data curation and formal analysis, reviewing and editing

**SP:** data curation and formal analysis, reviewing and editing

**GN:** investigation

**AL:** investigation

**SD:** investigation

**EDA:** investigation

**FG:** investigation

**GD:** reviewing and editing

**PTJS:** reviewing and editing, supervision, project administration and resources for study conduct.

**DM** conceptualization, investigation, visualization, writing the original work, reviewing and editing, supervision, project administration and resources for study conduct.

## Data availability

No datasets were generated or analysed during the current study

